# Pharmacological rescue of motor circuit dysfunction in a *Drosophila* model of paroxysmal dyskinesia

**DOI:** 10.1101/2024.07.05.602188

**Authors:** Abigail D. Wilson, Yuyao Jiang, Nidhika Desai, Nell Simon-Batsford, Angelina Sanderson, Isaac Tolley, Hao Gao, James E.C. Jepson

**Affiliations:** Department of Clinical and Experimental Epilepsy, UCL Queen Square Institute of Neurology, University College London, London, WC1N 3BG, UK; Champalimaud Research, Champalimaud Centre for the Unknown, Lisboa, Portugal

## Abstract

**Background:** Paroxysmal dyskinesias (PxDs) are characterised by bouts of involuntary dystonic and choreiform movements. The patho-mechanisms underlying these debilitating disorders remain poorly understood, and drug treatments are often limited. We recently generated a *Drosophila* model of a PxD-linked mutation causing BK potassium channel gain-of-function (BK GOF), and showed that BK GOF perturbs movement in *Drosophila* by disrupting neurodevelopment. However, whether locomotor capacity in BK GOF flies can be pharmacologically restored following neurodevelopmental insults has remained unclear.

**Objective:** To identify pharmacological suppressors of motor defects caused by BK GOF.

**Methods:** Using adult BK GOF flies, we performed an unbiased, in vivo, locomotion-based screen of 370 FDA-approved drugs. To test the impact of positive hits from this screen on motor circuit activity, we used optical imaging to record the intrinsic rhythmic activity of *Drosophila* larval motor circuits driving peristalsis and turning behaviors.

**Results:** We found that inhibitors of acetylcholinesterase – a protein that degrades acetylcholine in cholinergic synapses – partially rescued movement defects caused by BK GOF. Inhibition of acetylcholinesterase also partially restored intrinsic activity of motor circuits controlling forward movement and turning in BK GOF larvae.

**Conclusions:** Our findings indicate that elevating cholinergic tone can reverse motor circuit dysfunction in an animal model of PxD caused by BK GOF. Furthermore, our study provides proof-of-principle that *Drosophila* can be utilised for screens to uncover putative drug treatments for involuntary movement disorders.

## INTRODUCTION

Paroxysmal dyskinesias (PxDs) are debilitating disorders characterized by episodes of painful, involuntary movements. These movements can manifest as dystonia, which involves sustained or intermittent muscle contractions, and/or chorea, which consists of brief, irregular muscle contractions ^1,2^. PxDs can be clinically sub-divided based on the trigger for involuntary movements ^2^. One such subtype is paroxysmal non-kinesigenic dyskinesia (PNKD), in which caffeine, alcohol, stress, and fatigue, frequently act as triggers for dystonic and choreiform movements. Mutations in a number of presynaptic proteins and ion channels have been shown to cause PNKD, suggesting that PNKD represents a synaptopathy/channelopathy in which involuntary movements arise due to pathological changes in neurotransmitter release, synaptic plasticity, or neural excitability, across motor circuits ^1^. However, the critical patho- mechanisms that drive involuntary movements in PNKD remain obscure, and pharmacological therapies capable of counteracting motor circuit dysfunction in PNKD are limited.

Here we focus on a form of PNKD caused by gain-of-function (GOF) mutations in *KCNMA1*, which encodes the pore-forming α-subunit of the Ca^2+^ and voltage-activated large-conductance K^+^ (BK) channel ^3–6^. BK channels are broadly expressed throughout metazoan nervous systems and influence neural excitability by modulating action potential shape and pre-synaptic neurotransmitter release ^7,8^. BK GOF mutations are associated with PNKD, seizures and, in some cases, hypotonia, neurodevelopmental delay, intellectual disability and autism ^4^. The first BK GOF mutation identified – a dominant D434G mis-sense mutation – was found to increase open probability and Ca^2+^-sensitivity of the channel, and patients harboring this allele exhibit PNKD, tonic-clonic and absence seizures, or both ^3,9^. Recently, fruit fly (*Drosophila*) and mouse models of this mutation have been characterized, yielding insights into disease patho-mechanisms ^10–12^. BK GOF knock-in adult *Drosophila* exhibit locomotor defects, altered limb kinematics, and repetitive limb movements ^12,13^. During the larval stage of the *Drosophila* life cycle, BK GOF suppresses the intrinsic rhythmogenic activity of motor circuits in the ventral nerve cord (VNC, a region functionally analogous to the mammalian spinal cord ^14^) that drive forward/backwards peristalsis and turning during foraging ^12^. Hence, BK GOF perturbs both movement and the patterned activity of motor circuits in *Drosophila*.

BK GOF knock-in mice similarly display impaired locomotion ^10^, as well as ethanol-induced involuntary movements ^10^, hypotonia-like phenotypes ^11^, and hyperexcitability of cerebellar Purkinje cells coupled with alterations in somatic and dendritic morphology ^10^. Consistent with the neurodevelopmental morbidities observed in some BK GOF patients ^5,6^ and altered Purkinje cell development in BK GOF mice ^10^, we recently found that GOF BK channels specifically act during late neurodevelopment to perturb movement and limb kinematics in adult *Drosophila*, and impair synaptic maturation during this critical developmental period ^13^. These data suggest that BK GOF may cause PNKD by altering the development of motor circuits. Hence, drug treatments for BK GOF patients may need to ameliorate a ‘dysfunctional motor circuit end-state’ that arises following prior neurodevelopmental perturbations, rather than modulating BK channels directly ^13^. Indeed, some PNKD patients with BK GOF mutations respond well to amphetamine-based stimulants ^5,15^, which do not target BK channels. However, amphetamines also yield significant side effects, including insomnia and anorexia. Identifying additional pharmacological suppressors of motor dysfunction in BK GOF animal models may thus identify novel therapies for PNKD and reveal key aspects of motor circuit dysfunction emerging from neurodevelopmental insults caused by BK GOF.

To achieve this goal, we implemented an unbiased movement-based drug screen utilizing adult BK GOF fruit flies. From a library of 370 FDA-approved drugs we identified rivastigmine, an acetylcholinesterase (AChE) inhibitor, as an acute suppressor of locomotor defects in BK GOF flies. Rivastigmine also partially restored the intrinsic rhythmogenic activity of VNC motor circuits in *Drosophila* larvae. Since AChE inhibition elevates acetylcholine levels at cholinergic synapses, our data reveals cholinergic tone as an important modulator of motor dysfunction in this form of PNKD.

## RESULTS

### Pharmacological suppression of motor defects in BK GOF flies

To identify cellular pathways that could be modulated to rescue motor dysfunction caused by BK GOF, we screened for compounds that could acutely enhance locomotor capacity in BK GOF flies. The dominant *slowpoke* E366G mutation ^12^ (*slo*^E366G^, orthologous to *KCNMA1*^D434G^) was placed in trans with the TM6B balancer chromosome to generate a stable heterozygous stock of BK GOF flies. Female flies from this stock were fed fly food containing either a control vehicle (DMSO) or one of 370 FDA-approved drugs dissolved in DMSO (Figure 1A and Table S1; see Methods). Locomotor activity of flies was measured using the *Drosophila* Activity Monitor, which quantifies infra-red beam crosses made as flies traverse a glass tube ^16^. Each tube also contained corresponding vehicle or drugs dissolved in fly food.

**Figure 1.**
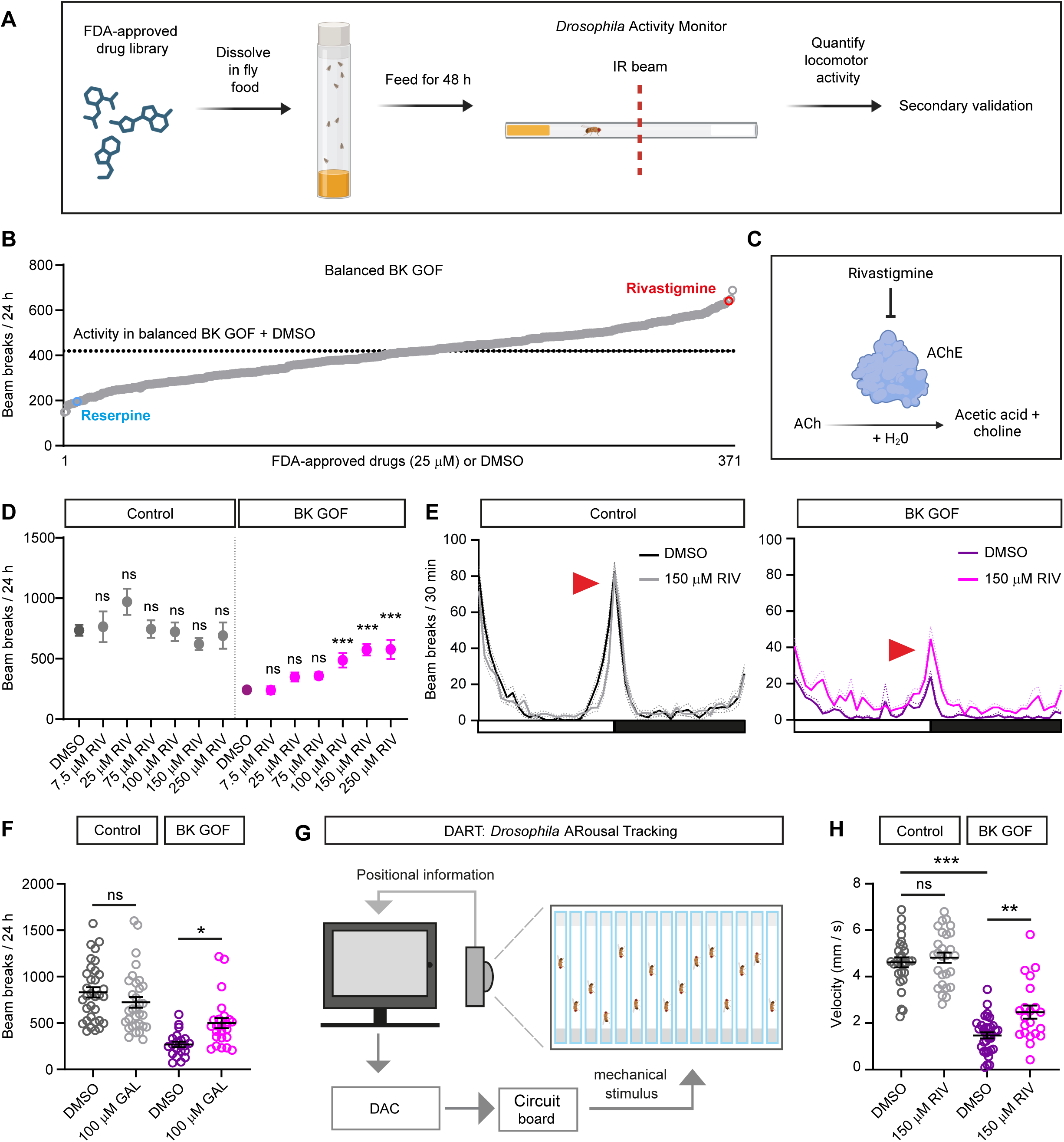
**Inhibitors of acetylcholinesterase partially rescue motor defects in BK GOF adult flies**. **A.** Schematic illustrating the screening protocol to identify pharmacological suppressors of movement defects in BK GOF adult flies. **B.** Locomotor activity in adult male balanced BK GOF flies (with the *slo*^E366G^ allele placed in trans with the TM6B balancer chromosome) fed 370 FDA-approved drugs (at 25 µM). Resulting overall movement (beam breaks in 24 h), quantified by the *Drosophila* Activity Monitor, is rank ordered. Effect of rivastigmine is noted in red. Effect of reserpine is noted in blue. Mean activity in *slo*^E366G^/TM6B flies fed a control vehicle (DMSO) is noted by the dashed line. n ≥ 5 adult male flies per drug (see Table S1). **C.** Schematic illustrating mode of action of rivastigmine. **D.** Rivastigmine (RIV) enhances overall locomotion (beam breaks per 24 h) in non-balanced BK GOF (*slo*^E366G/+^) adult male flies in a dose-dependent manner, but not in controls (*slo*^loxP/+^). Note that higher doses of rivastigmine are required to enhance movement in *slo*^E366G/+^ males compared to *slo*^E366G^/TM6B females (Figure 1B). Control: n = 8-69. BK GOF: n = 10-60. **E.** Mean locomotor activity (beam breaks per 30 mins) across 12 h light: 12 h dark periods in control and BK GOF adult males fed vehicle (DMSO) or 150 µM rivastigmine. *slo*^loxP/+^ + DMSO: n = 69, *slo*^loxP/+^ + 150 µM rivastigmine: n = 35, *slo*^E366G/+^ + DMSO: n = 60, *slo*^E366G/+^ + 150 µM rivastigmine: n = 36. Arrow indicates period of peak activity at lights off (ZT12). **F.** Schematic illustrating set-up of the *Drosophila* ARousal Tracking (DART) system. **G.** Peak locomotor velocity in control (*slo*^loxP/+^) and BK GOF (*slo*^E366G/+^) adult male flies fed DMSO or 150 µM rivastigmine, following DART-delivered mechanical stimuli. Left to right, n = 27, 27, 34, and 21. **H.** Overall locomotion (beam breaks per 24 h in *Drosophila* Activity Monitor) in control (*slo*^loxP/+^) and BK GOF (*slo*^E366G/+^) adult male flies fed vehicle (water) or 100 µM galantamine. Left to right, n = 33, 32, 21, and 24. Mean values are shown. Error bars: standard error of the mean (SEM). *p<0.05, **p<0.005, *** p<0.0005, ns – p>0.05, Kruskal-Wallis test with Dunn’s post-hoc test (D, I) or one-way ANOVA with Tukey’s multiple comparisons test (H).

Stimulants such as dextroamphetamine have previously been shown to improve motor phenotypes in BK GOF patients ^5,15^. Amphetamine enhances extracellular dopamine levels through a mechanism involving the dopamine transporter and vesicular monoamine transporters (VMATs) ^17^. Consistent with these observations, we found that a drug with opposing effects on dopamine levels – the VMAT inhibitor reserpine – enhanced the severity of motor defects in BK GOF flies (Figure 1B), an effect that was genotype-specific (Figure S1A). These data suggest that similar neuronal pathways can influence motor dysfunction in BK GOF humans and flies.

We therefore next sought to define compounds capable of specifically rescuing motor defects in the BK GOF background. Of the 370 drugs tested, 6 increased overall locomotion by > 50% compared to vehicle-treated flies (Figure 1B and Table S1). By re-testing these in control (*slo*^loxP/+^, see Methods) and non-balanced BK GOF (*slo*^E366G/+^) male flies, we identified rivastigmine as a specific and dose-dependent suppressor of motor defects caused by BK GOF in *Drosophila* (Figure 1B-E). Rivastigmine is a reversible non-competitive inhibitor of AChE, elevating synaptic ACh by preventing its hydrolysis in synaptic clefts (Figure 1C) ^18^. Acute feeding of rivastigmine (100-250 µM) increased both total locomotion over 24 h, and peak levels of self-driven locomotor activity, in BK GOF but not control male flies (Figure 1D-E and Figure S1B). Similar specific effects were observed in adult female flies (Figure S1C). We also tested whether galantamine, a distinct compound that also inhibits AChE, was similarly capable of increasing spontaneous locomotion in BK GOF in male flies. Indeed, acute feeding of 100 µM galantamine specifically enhanced overall locomotor activity in BK GOF but not control adult flies (Figure 1F), confirming that increasing acetylcholine levels through AChE inhibition enhances locomotor activity in BK GOF *Drosophila*.

However, we noted that rivastigmine appeared to elevate the activity of BK GOF flies across a 12 h light: 12 h dark period (Figure 1F), raising the possibility that rivastigmine might enhance overall activity through indirect effects (for example, via reduced sleep) instead of by improving locomotor capacity. To examine this more explicitly, we use the *Drosophila* Arousal Tracking (DART) system ^19^, which enables hi-resolution motion tracking through videography. We took advantage of DART’s ability to deliver mechanical stimuli to control and BK GOF flies, and quantified the velocity of the resulting startle responses (Figure 1G). Acute feeding of rivastigmine specifically enhanced stimulus-induced locomotor velocity in BK GOF adult male flies (Figure 1H), demonstrating an impact of rivastigmine on movement that is independent from confounding variables such as sleep and circadian rhythms.

### Rivastigmine restores rhythmogenic motor circuit activity in BK GOF larvae

Motor circuits responsible for patterning limb movements in *Drosophila* reside in a neuropil domain termed the ventral nerve cord (VNC) ^14^. We therefore sought to examine whether rivastigmine could restore functionality to VNC motor circuits in BK GOF *Drosophila*. To do so, we turned to *Drosophila* larvae for further functional analyses. The larval VNC contains rhythmogenic circuits that control aspects of larval locomotion such as forward and backward peristalsis, and turning/head-sweeping (which enables goal-oriented navigation during foraging) ^20,21^. In contrast to their adult counterpart, isolated larval brains exhibit intrinsic rhythmic activity of VNC motor circuits, manifesting as waves of motoneuron excitation spreading from posterior to anterior segments (fictive forward peristalsis), anterior to posterior segments (fictive backwards peristalsis), or unilateral excitation in anterior segments (fictive turns or head-sweeps, which we collectively refer to as ‘turns’) ^22,23^. Importantly, we previously showed that both fictive peristalsis and turning are strongly reduced in BK GOF *Drosophila* larvae ^12^.

Hence, in control and BK GOF larvae, we expressed GCaMP6m – a fluorescent reporter of intracellular calcium – in larval motoneurons (Figure 2A). We acutely fed larvae DMSO or rivastigmine (150 µM), isolated brains of resulting wandering L3 larvae 48 h later, and optically recorded input from rhythmogenic motor circuits to motoneuron dendrites (Figure 2B-J). These experiments revealed a striking genotype-specific effect of rivastigmine on rhythmic activity of larval VNC motor circuits. Acute rivastigmine feeding in control larvae was clearly detrimental to the activity of VNC motor circuits, partially reducing the proportion of isolated brains exhibiting rhythmogenic input to motoneurons reducing the frequency and amplitude of fictive forward waves (Figure S2A, B) and the number of fictive turns (Figure 2C- F, L). In contrast, in BK GOF larvae, rivastigmine feeding increased the proportion of isolated brains exhibiting rhythmogenic activity (Figure 2G-K) and the number of fictive turns (Figure 2L), without altering the frequency or amplitude of fictive forward waves (Figure S2A, B).

**Figure 2.**
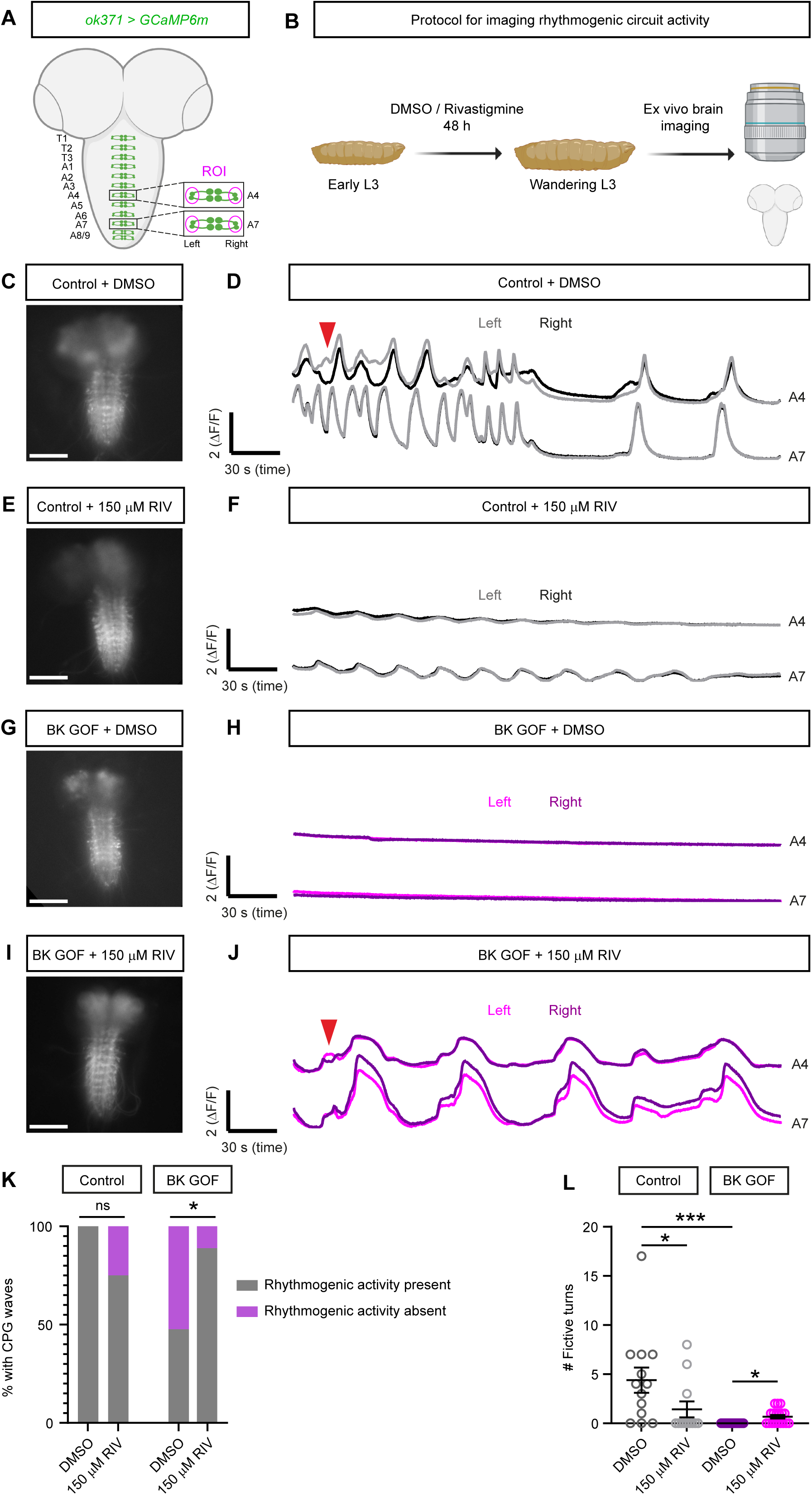
Acetylcholinesterase inhibition partially restores rhythmogenic motor circuit activity in BK GOF larvae. **A.** Schematic illustrating location of motoneuron cell bodies and dendritic clusters in the 3^rd^ instar larval *Drosophila* brain, labelled with GCaMP6m (*ok371 > GCaMP6m*). An example optical region of interest (ROI) encompassing lateral motoneuron dendritic clusters within abdominal segment 4 (A4) is shown. Location of the nine abdominal (A1-A8/9) and three thoracic segments (T1-T3) are also shown. **B.** Schematic showing larval drug feeding and imaging protocol. **C-F.** Epi-fluorescence images (C, E) and traces showing fictive peristaltic waves between segments A4 and A7 and fictive turning behavior (D, F) in control larvae fed DMSO or 150 µM rivastigmine. Red arrow (D) illustrates an instance of a fictive turn, characterised by asymmetric excitation in the anterior but not posterior abdominal segments. Scale bars: 100 µm. **G-J.** Epi-fluorescence images (C, E), and traces showing fictive peristaltic waves between segments A4 and A7 and fictive turning behavior (D, F), in BK GOF larvae fed DMSO or 150 µM rivastigmine. Red arrow (J) illustrates an instance of a fictive turn. Scale bars: 100 µm. **K.** Contingency table showing % of ex vivo larval brains exhibiting or not exhibiting intrinsic rhythmic input to motoneuron dendrites, from control or BK GOF larvae fed DMSO or 150 µM rivastigmine. Left to right, n = 13, 12, 21, and 18. **L.** Number of fictive turns in control or BK GOF larvae fed DMSO or 150 µM rivastigmine. Left to right, n = 13, 12, 21, and 18. Mean values are shown. Error bars: standard error of the mean (SEM). *p< 0.05, *** p< 0.0005, ns – p> 0.05, Fisher’s exact test (K) or Kruskal-Wallis test with Dunn’s post-hoc test (L).

Hence, rivastigmine primarily restores the ability to initiate activation of motor circuits driving forwards, backwards, and turning motions, in BK GOF larvae.

## DISCUSSION

Prior studies have used to *Drosophila* to identify pharmacological suppressors of seizure-like phenotypes, lethality, and motor defects, in models of epilepsy, neurodevelopmental disorders, and Parkinson’s disease ^24–27^. Here we show that *Drosophila* can also be used to uncover putative drugs for involuntary movement disorders. Prior screens for such compounds have used cellular alterations as core phenotypic readouts – for example, the mis-localisation of a mutant TorsinA protein linked to dystonia ^28^, or the capacity to process mis-folded proteins in the ER following expression of this protein ^29^. Deploying *Drosophila* has allowed us to take a complementary approach, using movement defects as a disease-relevant phenotype of interest for an in vivo screen.

Our results demonstrate that pharmacologically elevating cholinergic tone through AChE inhibition can partially rescue the effects of BK GOF on movement, of which observations from BK GOF patients ^5,6^, as well as rodent ^10^ and *Drosophila* models ^13^, suggest arise (at least in part) due to perturbations in motor circuit development. These findings are surprising, since numerous lines of evidence suggest that enhanced cholinergic signalling contributes to the pathology of dystonia, a core feature of the involuntary movements observed in BK GOF patients ^3,5,6^. For example, striatal cholinergic interneurons in rodent models of DYT1-*TOR1A* dystonia are paradoxically excited by D2-type dopamine receptor agonists, leading to elevated ACh release that perturbs synaptic plasticity at cortico-striatal synapses and potentially motor learning ^30,31^. Furthermore, trihexyphenidyl – a muscarinic antagonist – is an effective treatment for some dystonia patients ^32^, and conversely, cholinergic agonists can induce dystonia in humans and primates ^33,34^. Indeed, a small number of case-studies suggest that rivastigmine itself can induce dystonia in humans ^35,36^.

Consistent with these observations, our work suggests that AChE inhibition is detrimental to motor circuit function in a wild-type background, particularly in the context of the larval motor system. Here, acute feeding of rivastigmine suppressed intrinsic rhythmic activity of pre-motor circuits controlling forward and turning movements in wild-type larvae. Strikingly, however, the opposite was true in BK GOF larvae. Hence, we postulate that in the altered context of a nervous system expressing GOF BK channels, AChE inhibition may paradoxically promote motor circuit dysfunction.

How might such an effect occur? In insects, acetylcholine is the primary excitatory neurotransmitter, acting via post-synaptic nicotinic acetylcholine receptors (AChRs) ^37^. Muscarinic AChRs – which couple to excitatory or inhibitory pathways ^38^ – also play key roles in insects in regulating synaptic plasticity and the activity of rhythmogenic motor circuits influencing movement ^39–41^. Intriguingly, antagonism of muscarinic AChRs in wild-type ex vivo larval brains strongly phenocopies the effect of BK GOF on intrinsic motor circuit activity, reducing the frequency of fictive peristalsis and turns/head-sweeps ^41^. Hence, we speculate that neurodevelopmental alterations caused by BK GOF may disrupt signalling at muscarinic synapses, an effect that can be partially reversed by rivastigmine. Further experiments are required to test this hypothesis, and critically, to examine whether rivastigmine or other AChE inhibitors can ameliorate motor defects in vertebrate models of PNKD ^10,11^.

Overall, applying the methodologies described here to other *Drosophila* models of diseases characterised by involuntary movements ^42–46^ may allow researchers to tailor drug treatments to genetically defined movement disorders, and potentially, to cluster distinct movement disorders based on differential drug sensitivity, thus revealing shared or distinct features of their underlying motor circuit pathology ^1,47,48^.

## METHODS

### High-throughput drug screening

Age-matched adult female BK GOF (s*lo*^E366G^*/*TM6B*) Drosophila* were selected for the drug screen. Heterozygosity for the sloE366G allele accurately models the dominant effects of the *KCNMA1*^D434G^ allele (which is heterozygotic in corresponding patients). We note that in *Drosophila*, homozygosity for s*lo*^E366G^ is lethal ^12^. Female flies were used for the screen to ensure ample drug consumption as female flies have a higher requirement for food intake. The TM6B balancer stock line was used for drug testing to allow for high-throughput screening. Drugs were obtained from the DiscoveryProbe™ FDA-approved Drug Library (APExBIO, L1021) and mixed into standard fly food to a final concentration of 25 µM with 0.25 % v/v DMSO or H_2_O. Approximately 12 flies were placed on drug containing food alongside the appropriate vehicle control for 48 h at a constant temperature (25°C) under 12 h: 12 h light-dark cycles (12L: 12D). Flies were subsequently placed in behavioral tubes with drug-containing fly food, and total locomotion was quantified over the next 12L: 12D cycle in the *Drosophila* Activity Monitor (DAM, Trikinetics inc., MA, USA). The DAM is a high-throughput automated behavioral assay that quantifies free-moving *Drosophila* behavior via the number of times the individual animal crosses the midpoint of a behavioral tube ^16^. Drugs that increased locomotion over 24 h were repeated in the same format. Male and female flies without the TM6B balancer (*slo*^loxP/+^ (control) or *slo*^E366G/+^ (BK GOF)) were subsequently tested to confirm drug hits and specificity.

### Adult *Drosophila* drug feeding and locomotor analysis

Rivastigmine (Cambridge Bioscience) was fed at increasing concentrations (7.5 – 150 µM) with a constant 0.25 % v/v DMSO to BK WT or BK GOF *Drosophila*. 150 µM rivastigmine was selected as the optimum dose for subsequent experiments. Galantamine (Merck) was fed at 100 µM for 48 h. Spontaneous total locomotion was quantified for 24 h in the DAM following 48 h acclimatisation under 12L: 12D at 25°C as described previously ^12^. Mechanical stimulus-driven locomotor activity was recorded using the *Drosophila* ARousal Tracking (DART, BFK Lab, UK) system ^19^, as described previously ^12^. Flies were de-winged (mechanically via forceps) for this assay, since the BK GOF flies have a downturned wing phenotype that meant they were susceptible to becoming stuck on their back following mechanical stimulus. Flies were only included in the analysis if quiescent (< 1 mm/s speed in the 5 mins prior to the mechanical stimulation). Stimulus locomotor activity was quantified as the speed of each fly in the first 1min bin after the stimulus.

### Larval *Drosophila* drug feeding

Larvae were removed from standard fly food by suspension in 20% sucrose, gently washed in distilled H_2_O, and placed on fly food containing 150 µM rivastigmine or DMSO control. Age-matched L3 larvae were selected for experiments 48 h later.

### Live imaging

For live fluorescence recordings of larval motor circuit activity, age-matched L3 larvae of *ok371*-Gal4, UAS-*GCaMP6m*, BK WT or BK GOF, were dissected to isolate the entire CNS (brain and VNC intact). Larvae were dissected and imaged in a recording saline ^49^ of (in mM): 135 NaCl, 5 KCl, 4 MgCl_2_, 2 CaCl_2_, 5 TES and 36 sucrose, pH 7.15 with NaOH, on poly-lysine coated glass dishes. Recordings were taken immediately on an Optical Imager (Photometrics Prime BSI CMOS Camera with CoolLED pE-800, Scientifica) at 100 ms intervals over 5 mins (3000 frames). Fluorescence was quantified at 4 ROIs (each of 187 µm^2^ in area) drawn around the dendritic regions of motoneurons in abdominal segments A7 and A4 on each side of the VNC (Figure 2A, with background fluorescence subtracted. Peaks in fluorescence indicated increased intracellular Ca^2+^. Forward waves were defined as concurrent contralateral peaks at motoneuron A7 prior to motoneuron A4, backward waves were defined as concurrent contralateral peaks at motoneuron A4 prior to motoneuron A7. Fictive turns were defined as asymmetric fluorescence at only one side of motoneuron A4 and may represent turning or head sweep behavior. Amplitude was quantified from the left A4 motoneuron during peak fluorescence during forward waves. Fluorescence is plotted as ΔF/F over 5 minutes, where F was defined as the minimum fluorescent value occurring over the 5 min period within the ROI, and ΔF represents the fold-change increase in fluorescence from the baseline F.

## ACKNOWLEDGEMENTS

We thank Dr. Gabriel Aughey and Dr. Simon Lowe for comments on the manuscript. This study was funded by a MRC Senior Non-Clinical Fellowship (MR/V03118X/1) to Prof. James Jepson. Schematics shown in Figures 1 and 2 were created with <u>Biorender.com</u>.

## AUTHOR CONTRIBUTIONS

A.D.W – Conceptualization, Methodology, Investigation, Writing - Original Draft, Writing – Review & Editing, Visualization, Funding acquisition. Y.J – Investigation. N.D – Investigation. N.S-B – Investigation. A.S – Investigation. I.T – Investigation. H.G – Investigation. J.E.C.J – Conceptualization, Visualization, Writing - Original Draft, Writing – Review & Editing, Supervision, Project administration, Funding acquisition.

---

**Figure S1.**
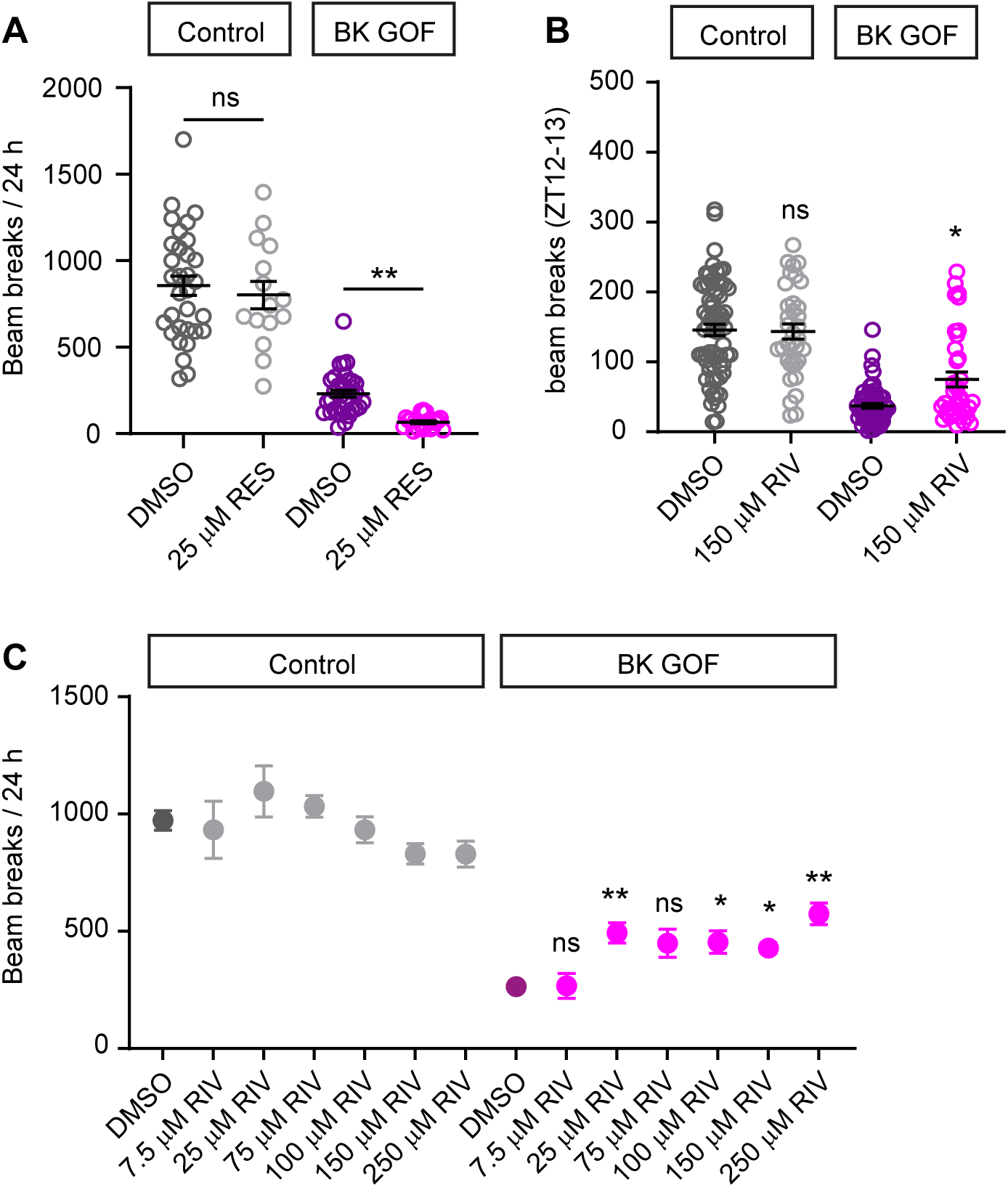
Opposing effects of reserpine and rivastigmine on locomotor capacity in adult BK GOF flies. A. Overall locomotion (beam breaks per 24 h in *Drosophila* Activity Monitor) in control (*slo*^loxP/+^) and BK GOF (*slo*^E366G/+^) adult male flies fed vehicle (DMSO) or 25 µM reserpine (RES). Left to right, n = 32, 15, 36, and 20. B. Rivastigmine (RIV) enhances peak locomotor activity (occurring in the one-hour period following lights-off in 12 h light: 12 h dark conditions) in non-balanced BK GOF (*slo*^E366G/+^) adult male flies. Left to right: n = 69, 35, 60, 36. C. Rivastigmine (RIV) enhances overall locomotion (beam breaks per 24 h) in BK GOF (*slo*^E366G/+^) adult female flies in a dose-dependent manner, but not in controls (*slo*^loxP/+^). control: n = 10-70. BK GOF: n = 9-80. Mean values are shown. Error bars: standard error of the mean (SEM). *p< 0.05, **p<0.005, ns – p> 0.05, Kruskal-Wallis test with Dunn’s post-hoc test.

**Figure S2.**
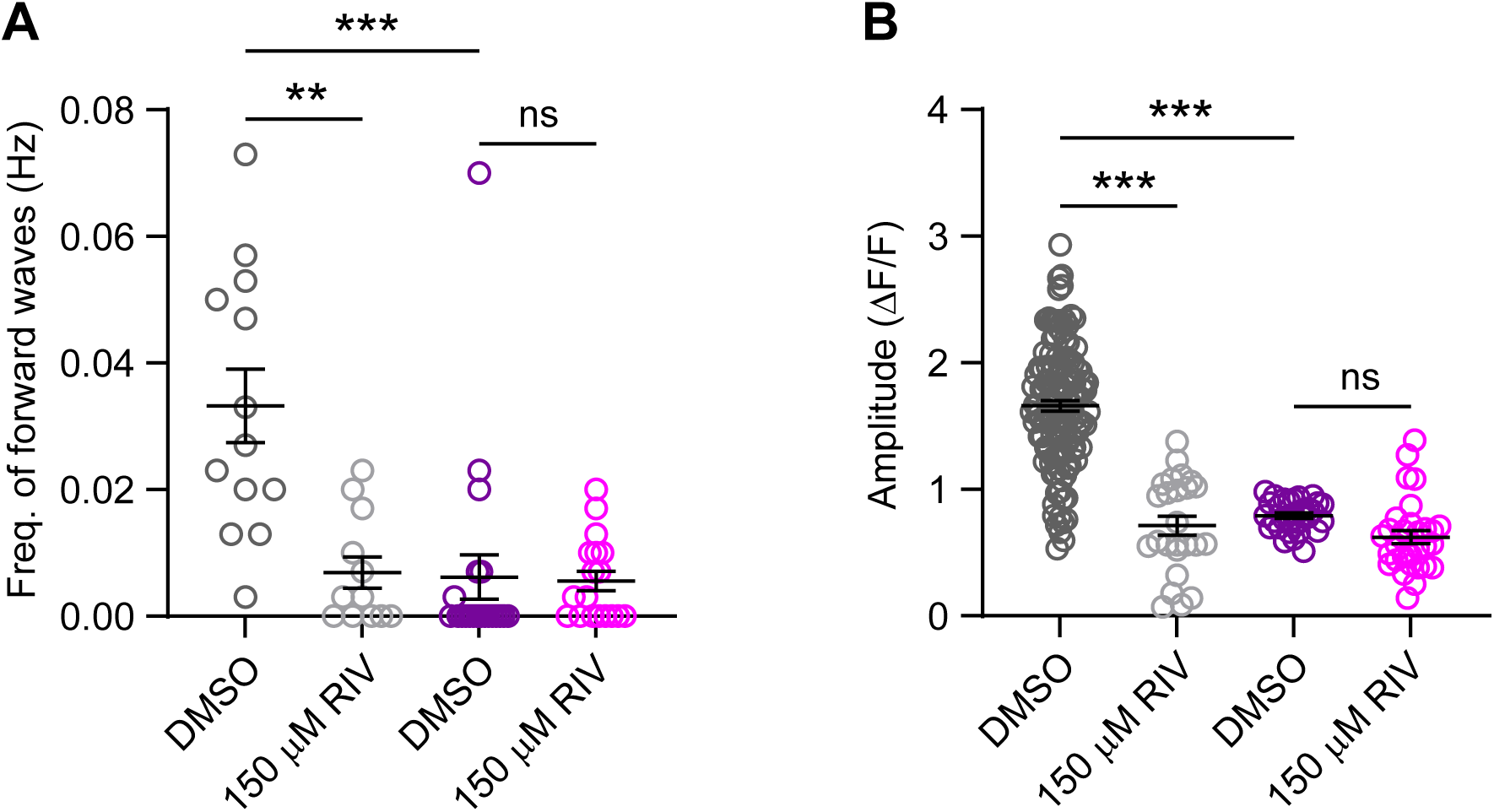
Rivastigmine does not rescue changes in the frequency or amplitude of forward fictive peristaltic waves in BK GOF *Drosophila* larvae. **A.** Frequency of forward fictive peristaltic waves in control or BK GOF larvae fed DMSO or 150 µM rivastigmine. Left to right, n = 13, 12, 21, and 18. **B.** Amplitude of forward fictive peristaltic waves in control or BK GOF larvae fed DMSO or 150 µM rivastigmine. Left to right, n = 30, 25, 31, and 30. Mean values are shown. Error bars: standard error of the mean (SEM). **p< 0.005, *** p< 0.0005, ns – p> 0.05, Kruskal-Wallis test with Dunn’s post-hoc test.

